# Active force generation shapes the metaphase spindle through a mechanical instability

**DOI:** 10.1101/2020.02.08.939868

**Authors:** David Oriola, Frank Jülicher, Jan Brugués

## Abstract

The metaphase spindle is a dynamic structure that segregates chromosomes during cell division. Recently, soft matter approaches have shown that the spindle behaves as an active liquid crystal. Still, it remains unclear how active force generation contributes to its characteristic spindle-like shape. Here, we combine theory and experiments to show that molecular motor driven forces shape the structure through a barreling-type instability. We test our physical model by titrating dynein activity in *Xenopus* egg extract spindles and quantifying the shape and microtubule orientation. We conclude that spindles are shaped by the interplay between surface tension, nematic elasticity and motor-driven active forces. Our study reveals how active force generation can mold liquid crystal droplets and it has implications on the morphology of non-membrane bound compartments demixed from the cytoplasm.

---

Cell division relies on the self-organization of the mitotic spindle, a highly dynamic non-equilibrium structure that segregates chromosomes to the two daughter cells. This structure is composed of an aligned array of microtubules – dynamic polymers that nucleate, grow and shrink – and energy transducing proteins such as molecular motors (*1*) that generate forces and fluxes within the structure. *In vitro* reconstitution approaches have advanced our basic understanding of how microtubule-motor mixtures self-organize (*2–6*). Motor activity can bundle, cluster and slide microtubules. These processes generate non-equilibrium patterns in bulk mixtures such as asters, vortices (*2, 6*) or even turbulent-like flows (*3*), that can be recapitulated by active liquid crystal descriptions (*7–9*). The self-organization of molecular motors and microtubules in spindles has been the subject of intense investigation (*10–13*). Nevertheless, how motor-driven forces determine the characteristic bipolar shape of the spindle remains poorly understood (*11, 14–17*).

Mitotic motors are known to critically influence spindle shape and microtubule orientation (*17–21*). Inhibition of dynein leads to unfocused poles and flag-like microtubule structures (see Fig. 1A) (*19, 22*), whereas inhibition of kinesin-5 leads to monopolar spindles (*17, 23*). How does the interplay of motor-induced forces and the liquid crystalline order govern spindle shape is unclear. Here we combine theory and experiments to understand how motor activity shapes *Xenopus* egg extract spindles. By considering the spindle as a liquid crystal droplet, we find that its characteristic bipolar shape emerges through a barreling-type instability as a consequence of dynein activity. The mechanism resembles the process by which an elastic beam acquires a barrel-like shape under the action of compressive forces (*24*). We experimentally test our theory by characterizing the evolution of the spindle shape and microtubule orientation under the titration of a dynein inhibitor. Overall, our results are consistent with spindle pole focusing being a consequence of a mechanical instability driven by dynein-mediated forces.

**Figure 1:**
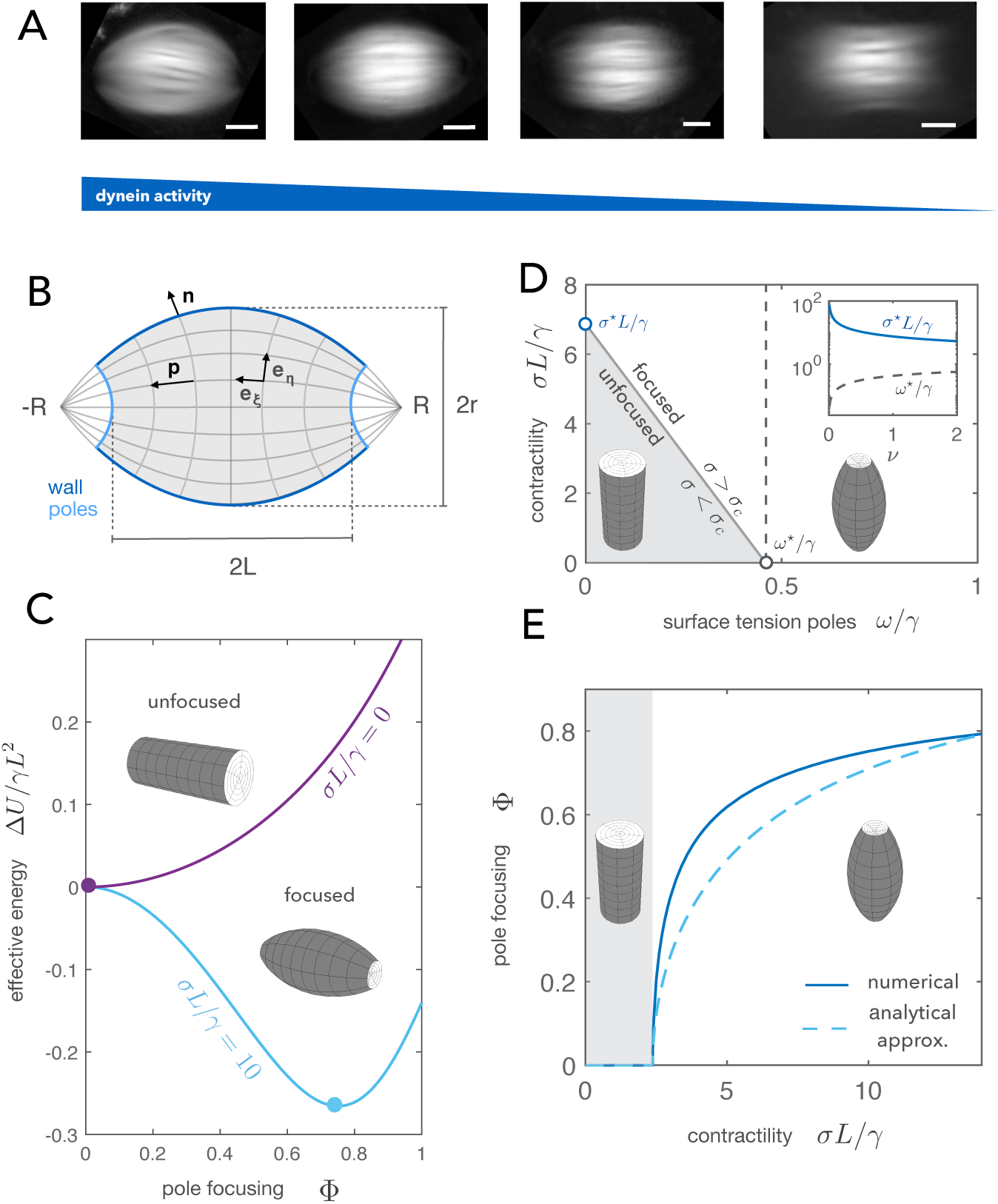
Theory of spindle pole focusing. A) Averaged retardance images of egg extract spindles under different concentrations of the dynein inhibitor p150-CC1. Images correspond to different titration days (Scale bars: 20 *µ*m). B) The spindle is described as a tactoid of length 2*L*, width 2*r* and pole-to-pole distance 2*R*. The two dimensionless parameters describing the spindle shape are the pole focusing parameter Φ = *L/R* and the aspect ratio *a* = *r/L*. The spindle is parametrised using bispherical coordinates {*ξ, η, φ*} and the director field **p** follows the *ξ*-coordinate (see SI). C) Dimensionless energy of the spindle relative to the cylindrical configuration Δ*U/γL*^2^ as a function of the pole focusing parameter Φ for two different values of the stress *σ*. The shape is found by minimising Δ*U* = *U*(Φ) − *U*(0). D) Phase diagram of how spindle shape changes as a function of the contractile stress *σ* and the surface tension at the poles *ω*. (Inset) Evolution of *σ*^⋆^ and *ω*^⋆^ as a function of the dimensionless volume *ν*. E) Bifurcation diagram considering dynein contractility *σ* as the control parameter: Below a certain critical contractility *σ*_*c*_ the cylindrical configuration Φ = 0 is stable. Once the contractility exceeds the threshold value *σ*_*c*_, the structure undergoes axisymmetric buckling acquiring a barrel-like shape. The solid curve corresponds to the full solution and the dashed curve corresponds to the analytical solution expanding the energy up to quartic order in Φ (see SI). The study is done in the limit of constant volume (*AL/γ* → ∞) with parameters *K*_1_*/K*_3_ = 1, *K*_1_*/γL* = 0.1, *ω/γ* = 0.3, *ν* = 1.2.

## Differential surface tension and bulk contractility control spindle pole focusing

To understand how motor forces lead to spindle pole focusing, we formulate a coarse-grained theory considering the spindle as a liquid crystal with director **p**, characterising the local axis of microtubule orientation, and microtubule density *ρ*. We consider the spindle as a nematic tactoid (*25–30*), which we describe using an orientational field **p** as depicted in Fig. 1B. For an equilibrium nematic liquid crystal, the bulk free energy *F*_*b*_ reads (*7, 9, 31*):

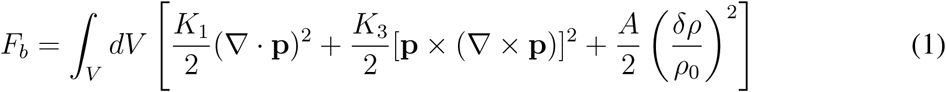

where *K*_1_, *K*_3_ are the Frank elastic constants (*31*), *A* is a compression modulus and *δρ* = *ρ* − *ρ*_0_ corresponds to density fluctuations around a preferred density *ρ*_0_ (*7*). The first two terms account for splay and bending deformations of a bulk nematic, respectively, while the last term penalizes density fluctuations. For simplicity, we do not include twist deformations or a saddle-splay term (*27*). At the interface with the surrounding cytoplasm, we adopt the following expression for the surface free energy *F*_*s*_ (*27*):

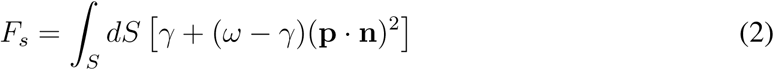

where **n** is the normal vector to the interface, *γ* is the surface tension and *ω* − *γ* is an anchoring strength describing how microtubules anchor to the spindle surface (*27, 30*). Motor activity on the surface can lead to active contributions to the surface tensions *γ* and *ω* (*32*). In addition to contributing to surface tension, motor proteins can also generate active stresses in the bulk (*7–9, 17*). These can be written as:

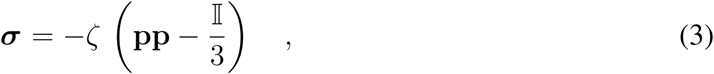

where the strength of the active stress is characterized by the coefficient *ζ*. The sign of *ζ* defines the extensile (*ζ* > 0) or contractile (*ζ* < 0) nature of the active stress (*7–9*). Motivated by *in vitro* studies, we consider the overall motor activity to be dominated by dynein and thus to be contractile (*4, 33, 34*). In general, active stresses can drive internal flows. In our approach, we neglect the role of viscous stresses arising from internal microtubule flows since they are not affected by dynein inhibition and the velocity gradients are small near the poles (see Fig. S1 and Materials and Methods). We parametrise the spindle by its length 2*L*, width 2*r* and the distance between the virtual poles 2*R* (see Fig. 1B). Such parametrisation allows us to continuously move from a tactoid to a cylinder, reproducing the shapes observed under the titration of dynein activity (see Fig. 1A). The spindle shape is determined by two parameters: the pole focusing parameter Φ = *L/R* and the aspect ratio *a* = *r/L*. Additionally, we consider the spindle length 2*L* to remain constant under changes in spindle shape. This constraint is motivated by recent evidence suggesting that spindle length is mainly set by microtubule nucleation gradients and tubulin mass balance but not by mechanics (*35, 36*).

Given the previous considerations, it can be shown that the curves *a*(Φ) are independent of the active stress (see SI). This allows us to reduce the problem to the minimisation of an effective work function *U* (Φ) that includes passive and active contributions. For simplicity, we consider the case where the spindle volume *V*_0_ is conserved and further show that compressibility effects do not significantly change the physics of the problem within the parameter regime relevant for the spindle (see Fig. S2 and SI). Close to the cylindrical configuration, the effective work function can be expanded up to fourth-order in Φ and the problem can be mapped to a Landau theory (*37*), where Φ plays the role of the order parameter:

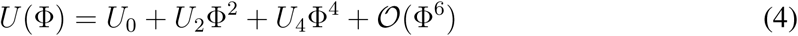

and the different coefficients can be explicitly determined (see SI). In particular, *U*_2_ changes sign as a function of the surface tension at the poles *ω* and the contractility parameter *σ* ≡ −2*ζ/*3 > 0. Provided that *U*_4_ > 0, axisymmetric buckling (also known as barreling (*24*)) occurs (see Fig. 1C,D,E) whenever *ω* or *σ* exceed a critical value and the cylindrical configuration (Φ = 0) becomes unstable (see Fig. 1D and Movie S1). Taking *σ* as our control parameter, the spindle undergoes a barreling transition at the critical stress (see Fig. 1E and SI):

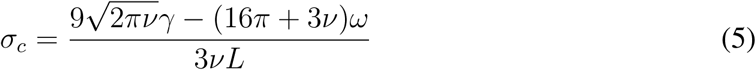

where *ν* = *V*_0_*/L*^3^. Our theory shows that contractile stresses as well as differential surface tension can focus spindle poles via a barreling-type instability. Contractility may focus the poles either by exerting normal stresses (*σ*) or in-plane stresses (*ω*) at the spindle poles, whereas nematic elasticity (*K*_1_, *K*_3_) and the surface tension at the spindle walls (*γ*) oppose to active forces.

## Dynein controls a barreling-type instability in spindles

Our theory predicts that spindle pole focusing is driven by a barreling-type instability. To verify this prediction, we experimentally quantified the changes in shape and microtubule orientation in spindles during the transition from closed to open poles by using an LC-Polscope (see Materials and Methods) and titrating the dynein inhibitor p150-CC1 (*22*). The transition was found to be reversible and poles focused by adding back fresh extract (see Materials and Methods and Fig. S3).

We characterised spindle shapes using the retardance images (Fig. 1A) to obtain the distance between the virtual poles (2*R*), spindle length (2*L*) and spindle width (2*r*) (see Materials and Methods and Fig. S4). The steady state spindle length and width were not significantly affected by the inhibition of dynein and read 2*L* = 57 ±14 *µ*m and 2*r* = 26 ± 9 *µ*m (*n* = 364, mean ± SD), respectively. Although spindles elongated right after the addition of p150-CC1, their steady state length after ∼ 30 min of inhibitor addition was not significantly affected during the titration (see Fig. S5, left), consistent with microtubule nucleation mainly determining spindle size (*35,36*). At the same time, the steady state spindle volume remained constant (see Fig. S5, right) in agreement with previous studies where spindle volume recovers after external deformations (*38*). The evolution of the microtubule orientational field was quantified using the slow axis component of the LC-Polscope (Fig. 2A), and showed good agreement to the prescribed orientational field (Fig. 2B,C). This result is a strong evidence that microtubule orientation in spindles is determined by passive liquid crystal elasticity and validates our choice of orientational field (*39*).

**Figure 2:**
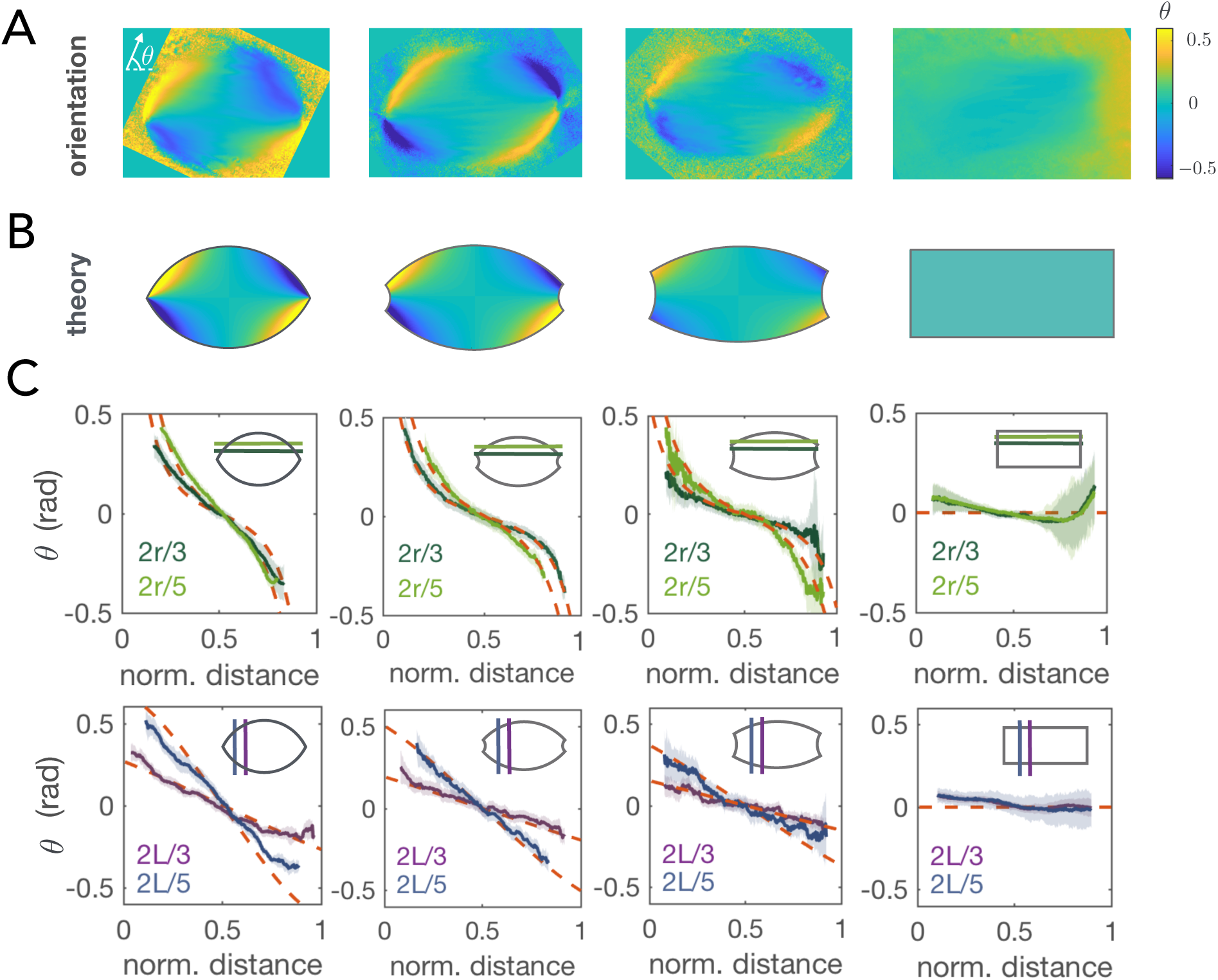
The microtubule orientational field in the spindle corresponds to a bispherical orientational field. A) Averaged orientational fields corresponding to the different spindles in Fig. 1A where *θ* is the orientation angle respect to the spindle long-axis. B) Prescribed orientational field **p** = **e**_*ξ*_*/* |**e**_*ξ*_| according to the shape parameters measured using the retardance signal. The angle range is chosen for visualisation purposes. C) Comparison between the prescribed orientational field in B (red dashed lines) and the averaged experimental orientational field in A (solid lines) for different sections along the spindle long axis (upper panel) and along the spindle short axis (lower panel). The shaded error bars correspond to SD.

Finally, for each titration we plotted Φ versus the rescaled inhibitor concentrations (see Materials and Methods) (Fig. 3). By approximating the active stress as a linear decreasing function of the inhibitor concentration (see caption Fig. 3) and estimating the parameters of the model (see SI and Table 1) we found good agreement between experiments and theory with no need of any fitting parameters (Fig. 3). Thus, we conclude that a barreling-type instability captures the process of pole focusing in spindles.

**Figure 3:**
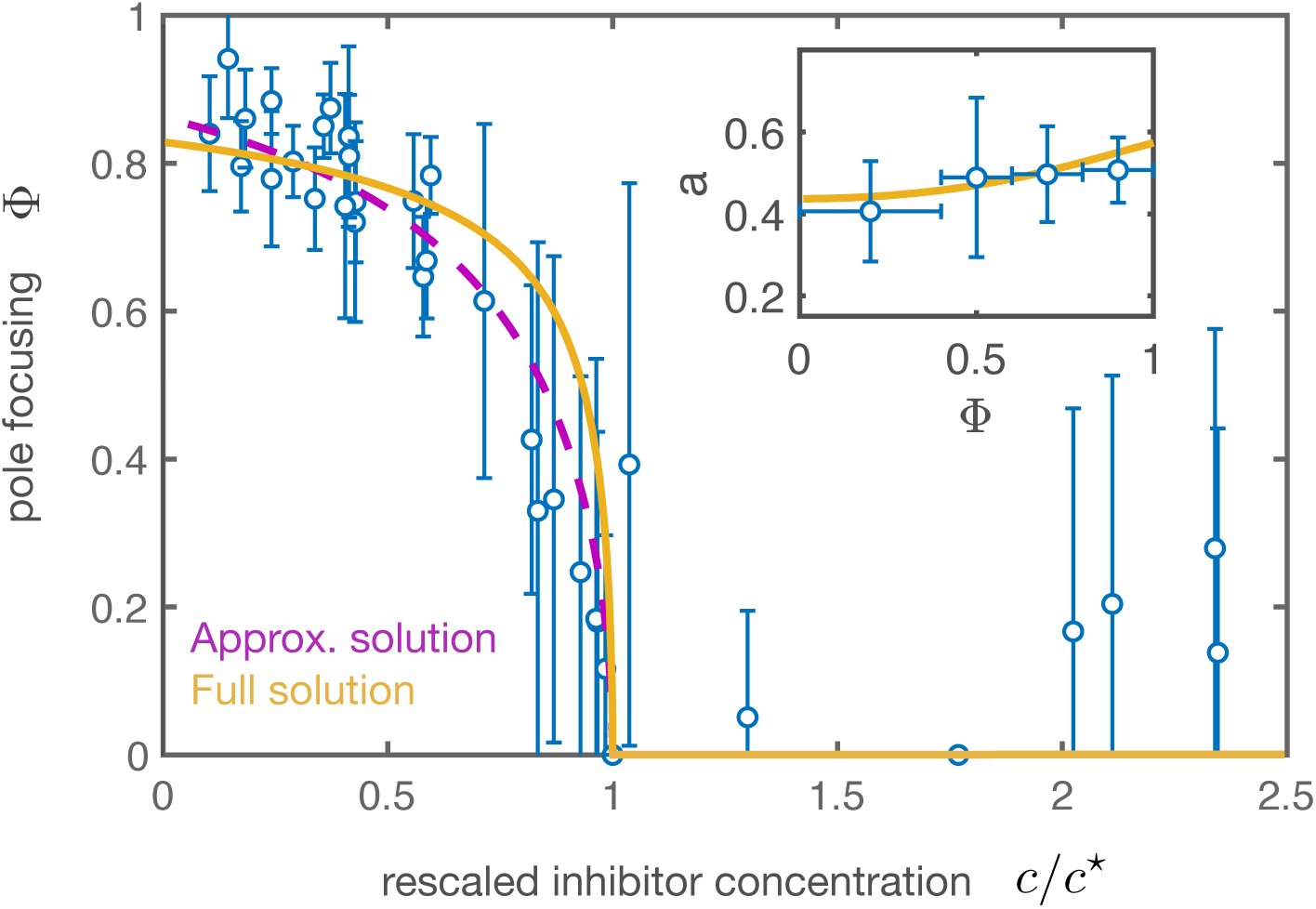
Bifurcation diagram under the titration of a dynein inhibitor. Pole focusing parameter Φ as a function of the rescaled concentration of p150-CC1 inhibitor (*c/c*^⋆^) with respect to the critical inhibitor concentration *c*^⋆^ for the corresponding extract day. The circles correspond to the averaged experimental results using data from 10 different titrations during 7 different extract days (*n* = 364 spindles). Error bars indicate SD. The purple dashed curve and the yellow solid curve correspond to the dashed and solid curves in Fig. 1E, respectively, using *c/c*^⋆^ = 1 − (*σ* − *σ*_*c*_)*/*(*σ*_max_ − *σ*_*c*_), where *σ*_max_ is the dynein stress at zero inhibitor concentration. *σ*_max_*L/γ* = 20, *σ*_*c*_*L/γ* = 2.376. The vertical error bars denote SD. (inset) Aspect ratio a as a function of the pole focusing parameter Φ (mean ± SD, *n* = 364 spindles). The theoretical curve (yellow) stems from the constant volume condition and has no fitting parameters (see SI). The parameters used for the theoretical curves are the same as in Fig. 1.

## Discussion

We have shown that pole focusing in spindles is a consequence of a mechanical instability driven by dynein activity. Dynein has been reported to generate active contractile stresses in disordered networks as a consequence of its accumulation at microtubule minus-ends (*4, 33, 34*). Our results are consistent with dynein focusing spindle poles by either generating contractile stresses or increasing the surface tension at the poles, where dynein is known to be enriched (*40*).

Although we have shown that dynein drives pole focusing, it is less clear what drives pole unfocusing in the absence of dynein. Our model provides three different ways to unfocus the spindle poles: via active dipolar extensile stresses (*ζ* > 0), increasing nematic elasticity (*K*_1_, *K*_3_) or decreasing the surface tension at the poles (*ω*) (see Fig. 1D). A potential candidate to account for these effects is kinesin-5, a tetrameric plus-ended motor that generates poleward flows by sliding antiparallel microtubule overlaps (*40,41*). One possibility is that kinesin-5 motors generate extensile stresses in the spindle, similarly as artificial tetrameric motor clusters in *in vitro* active nematic systems (*3*). However, microtubule overlaps dramatically decrease close to the poles (*13*). Consistent with this picture, unfocusing of spindle poles can be achieved in the presence of a kinesin-5 inhibitor (FCPT), forcing tight binding of kinesin-5 onto microtubules and blocking its ATP activity (*42*). We thus propose that kinesin-5 might unfocus spindle poles by bundling microtubules in the spindle thus increasing nematic elasticity and/or decreasing the effective surface tension at the poles.

Our study also provides an explanation to the puzzling problem of a bipolar structure under the double inhibition of kinesin-5 and dynein motors (*13, 17, 43*). In this case, spindle shape is set by a combination of differential surface tension and nematic elasticity, which in general can lead to a tactoidal shape with partially focused poles (Fig. 1D, *σ* = 0). Kinesin-5 bundles microtubules, thus leading to a flag-like configuration of the spindle in the absence of dynein activity. When dynein is added to the system, it focuses the poles by overcoming the stresses generated by kinesin-5, thus resuming a spindle-like shape. This raises the question of why the competition between kinesin-5 and dynein motors is necessary to set spindle shape given that a spindle-like structure is achieved in the absence of the two types of motors. A possible answer is that kinesin-5 poleward flux is essential for the proper microtubule organization in *Xenopus* extract spindles (*13, 44, 45*). However, as a side product of this activity, kinesin-5 induces pole unfocusing in the absence of dynein. Thus, we propose that dynein activity is essential to maintain a spindle-like structure to overcome pole unfocusing driven by kinesin-5, in line with recent studies in mammalian spindles (*46*).

More generally, our work shows that the spindle behaves as an active nematic droplet, similarly to other biological condensates such as actin (*30, 47, 48*) or microtubule (*49*) tactodial droplets. Indeed, the spindle shares some signatures with liquid droplets such as fusion and rapid internal rearrangement (*15, 50*). Further work will be needed to elucidate to what extent the physics of active nematic droplets can be used to understand cell division (*47, 51, 52*).

## Materials and Methods

### Cytoplasmic extract preparation, spindle assembly and biochemical perturbations

Cytostatic factor (CSF)-arrested *Xenopus laevis* egg extract was prepared as described previously (*53,54*). In brief, unfertilized oocytes were dejellied and crushed by centrifugation. After adding protease inhibitors (LPC: Leupeptin, Pepstatin, Chymostatin) and Cytochalasin D (CyD) to a final concentration of 10 *µ*g/ml each to fresh extract, we cycled single reactions to interphase by adding frog sperm (to 300-1000 sperm/*µ*l final concentration) and calcium solution (10 mM CaCl_2_, 250 mM KCl, 2.5 mM MgCl_2_ to 0.4 mM Ca^++^ final concentration), with a subsequent incubation of 1.5 h. While fresh CSF extract containing LPC and CyD was kept on ice, all incubation steps were performed at 18-20 ^*°*^C. The reactions were driven back into metaphase by adding 1.3 volumes of fresh CSF extract (containing LPC and CyD). Spindles formed within 30 min of incubation. We inhibited dynein with p150-CC1, purified according to Ref. (*55*) and added to the reactions to the desired final concentration and incubated for an additional ∼ 20 min. Prior to imaging, Höchst 33342 was added to the reactions to a final concentration ∼ 16 *µ*g/ml, to visualize DNA. For each titration, four different concentrations of p150-CC1 were used in 30 *µ*L reactions using the same extract batch, in the range from 1 to 8 *µ*M. Backward titrations were set in parallel to the forward titrations and increasing amounts of crude metaphase extract were added after 30 min to the different dynein-inhibited reactions. Titrations were repeated for 8 different extract days. 6 *µ*L of the previous extract reactions were dropped on MatTek glass bottom dishes and covered with 1 mL mineral oil to prevent the evaporation of the drops. In these conditions, spindles were remarkably stable and could be imaged continuously for more than 30 min. The critical dynein inhibitor concentration showed variability depending on the extract day; therefore, we estimated the critical concentration for each day and normalized the inhibitor concentration with respect to this value.

### LC-Polscope

Spindles were imaged using an LC-Polscope (on a Ti Eclipse microscope body) with an sCMOS camera (Hamamatsu Orca Flash 4.0) using a 60x 1.2 NA water immersion objective. An open chamber was used to avoid possible mechanical stresses which could affect the degree of pole focusing. For data acquisition we used *µ*Manager (*56*). The exposure time was set to 200 ms. A total number of 364 spindles were analyzed. In order to estimate the shape parameters we followed a similar characterization as in Ref. (*29*) and fit two intersecting circles forming a tactoid to the spindle retardance images. The threshold retardance value to define the spindle length 2*L* was the background level for each spindle (see Fig. S4). Close to the completely unfocused configuration the position of the virtual poles was difficult to estimate, thus we considered a spindle to be unfocused below a threshold value of the pole focusing parameter of ∼ 0.4. The room temperature was kept at 19 ^*°*^C.

### Speckle microscopy

Speckle microscopy on spindles was done by adding Atto 565 frog tubulin in egg extracts to a final concentration of ∼ 1 nM and imaging using a Nikon spinning disk microscope (Ti Eclipse), an EMCCD camera (Andor iXon DU-888 or DU-897), a 60x 1.2 NA water immersion objective, and the software AndorIQ for image acquisition. The tubulin speckles were further analysed using TrackMate (*57*) and classified according to which pole they moved to. The speed and density of speckle tracks in each population was obtained and the center of mass velocity was computed.

## Authors’ contributions

D.O. performed theoretical and experimental research, data analysis and drafted the manuscript. J.B and F. J. directed the project and drafted the manuscript. D.O and J.B. conceived the project.

## Acknowledgements

We thank F. Decker, B. Dalton, F. Berndt, K. Ishihara, E. Rieckhoff and T. Quail for critical reading of the manuscript. We also thank K. Ishihara for providing the construct to purify the protein p150-CC1, and J. Sharpe and J. Baumgart for useful discussions. We acknowledge funding from EMBO (Long-Term Fellowship with number 483-2016 to D.O.) and HFSP (CDA 74/2014 to J.B.).

