## Supplementary Material for "Active force generation shapes the metaphase spindle through a mechanical instability"

### Liquid crystal theory for the metaphase spindle

#### Bispherical coordinates

Let us consider a point in the spindle as $\mathbf{X}(s^1, s^2, s^3)$ where $s^i$ are generalized coordinates. The metric is defined as $g_{ij} = \mathbf{e}_i \cdot \mathbf{e}_j$, where $\mathbf{e}_i = \partial_i \mathbf{X}$ with $\partial_i = \partial/\partial s^i$. We denote a differential of volume as $dV = \sqrt{g} ds^1 ds^2 ds^3$ with $g = \det g_{ij}$. Similarly, a surface element in the area spanned by $\{s^1, s^2\}$, reads $dS = \sqrt{g'} ds^1 ds^2$ where $g' = \det g_{ij}$ and $i = 1, 2$. We parametrize the spindle using bispherical coordinates $\{s^1, s^2, s^3\} = \{\xi, \eta, \varphi\}$ (*1*) and use the simple ansatz $\mathbf{p} = \mathbf{e}_\xi / |\mathbf{e}_\xi|$ for the orientational field as depicted in Fig. 1B. The transformation from cartesian to bispherical coordinates reads:

$$\mathbf{X} = (x, y, z) = R \left( \frac{\sin\xi \sin\eta \cos\varphi}{1 + \sin\xi \cos\eta}, \frac{\sin\xi \sin\eta \sin\varphi}{1 + \sin\xi \cos\eta}, \frac{\cos\xi}{1 + \sin\xi \cos\eta} \right) \tag{1}$$

where $z$ runs along the spindle long axis. The non-zero elements of the metric tensor read:

$$g_{\xi\xi} = \frac{R^2}{(1 + \cos\eta \sin\xi)^2}; \quad g_{\eta\eta} = \frac{R^2 \sin^2\xi}{(1 + \cos\eta \sin\xi)^2}; \quad g_{\varphi\varphi} = \frac{R^2 \sin^2\eta \sin^2\xi}{(1 + \cos\eta \sin\xi)^2} \tag{2}$$

The spindle shape can be varied by changing only two parameters: the spindle width $2r$ and pole-to-pole distance $2R$. The spindle length $2L$, is assumed to be constant and to be set by a microtubule nucleation mechanism that sets the size of the structure (*2*). In dimensionless form we have two free parameters: the aspect ratio $a = r/L$ and the pole focusing parameter $\Phi = L/R$. The last parameter allows us to move continuously from a spindle with completely focused poles to a cylindrical configuration with completely unfocused poles. The coordinate $\xi \in [\xi_0, \pi - \xi_0]$ moves along circular arcs that connect the two poles, where $\xi_0 = \arctan[1/(2\Phi) - \Phi/2]$. On the other hand, the coordinate $\eta \in [0, \eta_0]$ moves along circular arcs with the centers at the poles and $\eta_0 = 2\arctan(\mathrm{a}\Phi)$ denotes the position of the spindle walls (see Fig. 1B). Finally the azimuthal angle range is $\varphi \in [0, 2\pi]$.

#### Bulk free energy of the spindle

By using the prescribed shape and orientational field, we can compute $F_b$ and $F_s$ as a function of the shape parameters $a$ and $\Phi$. The total free energy can be computed by using Eqs. 1 and 2 from the Main Text:

$$F = F_s + F_b = \gamma S_w + 2\omega S_c + F_1 + F_3 + F_V \tag{3}$$

where $S_w$ is the surface of the spindle wall and $S_c$ is the surface of a spindle cap (see Fig. 1B). The different terms are obtained by computing the integrals:

$$S_c = 2\pi \int_0^{\eta_0} d\eta \sqrt{g_{\eta\eta}(\xi_0, \eta) g_{\varphi\varphi}(\xi_0, \eta)} \tag{4}$$

$$S_w = 2\pi \int_{\xi_0}^{\pi-\xi_0} d\xi \sqrt{g_{\xi\xi}(\xi, \eta_0) g_{\varphi\varphi}(\xi, \eta_0)} \tag{5}$$

$$F_1 = \frac{4\pi K_1}{R^2} \int_{\xi_0}^{\pi-\xi_0} d\xi \int_0^{\eta_0} d\eta \sqrt{g} \cot^2 \xi \tag{6}$$

$$F_3 = \frac{\pi K_3}{R^2} \int_{\xi_0}^{\pi-\xi_0} d\xi \int_0^{\eta_0} d\eta \sqrt{g} \sin^2 \eta \tag{7}$$

$$F_V = \frac{A}{2} \left(1 - \frac{V_0}{V}\right)^2 V \tag{8}$$

where the volume $V$ reads:

$$V = 2\pi \int_{\xi_0}^{\pi-\xi_0} d\xi \int_0^{\eta_0} d\eta \sqrt{g} \tag{9}$$

#### Variational of work of the active stresses

In contrast to passive nematic droplets (*1, 3*), the spindle can generate stresses on its own as a consequence of the action of motor proteins. In general, the virtual work generated by the active stresses might not have an associated work function. However, in our particular case, it is possible to integrate the virtual work and obtain such function. We consider a deformation of the tactoid by infinitesimally varying the pole-to-pole distance and the width, $R' = R + \delta R$,

$r' = r + \delta r$, respectively. From this, we define two independent small parameters $\epsilon \equiv \delta R/R$ and $\rho \equiv \delta r/r$. The deformation will lead to a transformation $\eta' = \eta'(\xi, \eta)$ and $\xi' = \xi'(\xi, \eta)$. During this deformation the points lying on the $\xi = \pi/2$ plane are deformed by a scaling factor $r'/r$, while the points along the line $\eta = 0$ are not deformed. This leads to a nonlinear transformation which to linear order in $\epsilon, \rho$ reads:

$$\xi'(\xi) \approx \xi + \epsilon \cos \xi \tag{10}$$

$$\eta'(\eta) \approx \eta - (\epsilon - \rho) \sin \eta \tag{11}$$

The variational $\delta X_i$ can be simply calculated as:

$$\delta X_i(\xi, \eta, \varphi; \epsilon, \rho) = X_i'(\xi', \eta', \varphi') - X_i(\xi, \eta, \varphi) \tag{12}$$

Given that the orientational field $\mathbf{p}$ is chosen to be tangential to the spindle walls, the only non-zero contribution of the active stress is found at the spindle caps. The variational of work generated by the active stress on the two caps will be:

$$\delta W = -2 \int_S \sigma_{ij} n_j \delta X_i dS \tag{13}$$

A general deformation of the surface can be expressed as $\delta \mathbf{X} = \delta X_n \mathbf{n} + \delta X_a \mathbf{e}_a$, where $a = \eta, \varphi$ and $\mathbf{n} = -\mathbf{e}_\xi/|\mathbf{e}_\xi|$. We notice that $\mathbf{p} = -\mathbf{n}$ at the caps, therefore the integrand in Eq. 13 reduces to $-2\sigma\delta X_n$, where $\sigma = -2\zeta/3$. By using the coordinates $s^1 = \eta$ and $s^2 = \varphi$ , the non-zero metric elements in a cap surface located at $\xi_0$ are given by $g_{\eta\eta}(\xi_0, \eta)$ and $g_{\varphi\varphi}(\xi_0, \eta)$, where $\xi_0 = \arctan[1/(2\Phi) - \Phi/2]$. Let us consider a small deformation of a cap surface such that the initial surface is $\mathcal{S} = \{X_i \,|\xi = \xi_0, \eta \in [0, \eta_0], \varphi \in [0, 2\pi)\}$ and the deformed surface is $\mathcal{S}' = \{X_i' \,|\xi' = \xi_1, \eta' \in [0, \eta_1], \varphi' \in [0, 2\pi)\}$, where $\xi_0 = \xi_0(L, R)$, $\eta_0 = \eta_0(L, R, r)$ and $\xi_1 = \xi'(\xi_0)$, $\eta_1 = \eta'(\eta_0)$, with $\eta_0 = 2\arctan(\mathrm{a}\Phi)$. After some algebra, it can be shown that the magnitude of the normal displacement reads:

$$\delta X_n(\eta, \Phi) = -2\epsilon L \left( \frac{\tan^2\left(\frac{\eta}{2}\right)}{1 + \Phi^2 \tan^2\left(\frac{\eta}{2}\right)} \right) \tag{14}$$

Eq. 13 can be rewritten as:

$$\delta W \;=\; -2\sigma \int_0^{2\pi} d\varphi \int_0^{\eta_0} d\eta \sqrt{g_{\eta\eta}(\xi_0,\eta) g_{\varphi\varphi}(\xi_0,\eta)}\,\delta X_n \tag{15}$$

Noticing that $\delta\Phi = -(L/R^2)\delta R = -\Phi\epsilon$, and after some additional algebra, we obtain:

$$\delta W(\Phi, a) = -2\pi L^3 a^4 \sigma\, \Phi \left(\frac{-1+\Phi^2}{1+a^2\Phi^4}\right)^2 \delta\Phi \tag{16}$$

#### Energy minimisation

From Eq. 16, we see that the active stresses do not affect the aspect ratio of the spindle ($\delta W/\delta a$ = 0), but only change the pole focusing parameter ($\delta W/\delta\Phi \neq 0$). Hence, we can find an implicit relationship $a(\Phi)$ by numerically solving $\partial_a F(a,\Phi) = 0$ (see Fig. S2) . By substituting the last expression into Eq. 16, the effective work function generated by the active stress reads:

$$W(\Phi) = \int_0^{\Phi} \frac{\delta W(\Phi', a(\Phi'))}{\delta \Phi'} d\Phi' \tag{17}$$

In order to solve the energetic problem we define the total effective work function as $U(\Phi) = F(\Phi, a(\Phi)) + W(\Phi)$ and minimise respect to $\Phi$ ($\partial_\Phi U = 0$) to find the equilibrium pole focusing parameter $\Phi_0$. The shape will be given by $\{\Phi_0, a(\Phi_0)\}$.

#### Study close to the transition point in the limit of constant volume

In the constant volume limit $(AL/\gamma \to \infty)$ the condition $\partial_a F(a,\Phi) = 0$ is equivalent to the condition $V_0 = V(a,\Phi)$. In this limit, there exist a single curve $a(\Phi)$ which only depends on $\nu$. In Fig. 3 (inset), $a(\Phi)$ is compared to the experimental curve considering the experimentally measured value of $\nu$. It is instructive to study this limit close to the transition where $\Phi \ll 1$. Up to quadratic order in $\Phi$, the volume of the spindle reads:

$$\nu \equiv \frac{V_0}{L^3} = 2a^2\pi + a^2\pi \left(a^2 - \frac{4}{3}\right)\Phi^2 + \mathcal{O}(\Phi^3) \tag{18}$$

Solving for the aspect ratio $a$, we obtain:

$$a(\Phi) = \sqrt{\frac{\nu}{2\pi}} + \nu^{1/2}\left(\frac{8\pi - 3\nu}{24\sqrt{2}\pi^{3/2}}\right)\Phi^2 + \nu^{1/2}\left(\frac{64\pi^2 - 80\pi\nu + 21\nu^2}{384\sqrt{2}\pi^{5/2}}\right)\Phi^4 + \mathcal{O}(\Phi^6) \quad (19)$$

Using this expression, we can find the effective work generated by the active stress:

$$W(\Phi) = \frac{\sigma\nu^2 L^3}{4\pi}\Phi^2 - \nu^2 L^3 \sigma\left(\frac{4\pi + 3\nu}{48\pi^2}\right)\Phi^4 + \mathcal{O}(\Phi^6) \quad (20)$$

The total effective work function can be expanded around $\Phi = 0$ and we obtain a Landau-type energy:

$$U(\Phi) = U_0 + U_2\Phi^2 + U_4\Phi^4 + \mathcal{O}(\Phi^6) \quad (21)$$

where the coefficients read:

$$U_0 = L^2(2\sqrt{2\pi\nu}\gamma + \omega\nu) \quad (22)$$

$$U_2 = \frac{\nu^2 L^3}{4\pi}(\sigma_c - \sigma) \quad (23)$$

$$U_4 = \frac{L\nu^{1/2}}{576\pi^2}\left[(b_1\gamma + b_2\omega + b_5\sigma L)L + b_3 K_3 + b_4 K_1\right] \quad (24)$$

and the coefficients $b_i$ read:

$$b_1 = \sqrt{2\pi}(64\pi^2 - 192\pi\nu - 153\nu^2) \quad (25)$$

$$b_2 = 72\nu^{5/2} - 288\pi\nu^{3/2} + 64\pi^2\nu^{1/2} \quad (26)$$

$$b_3 = 288\pi\nu^{3/2} \quad (27)$$

$$b_4 = 1536\pi^2\nu^{1/2} \quad (28)$$

$$b_5 = 12(4\pi + 3\nu)\nu^{3/2} \quad (29)$$

being $\sigma_c$ the critical contractile stress:

$$\sigma_c = \frac{9\sqrt{2\pi\nu}\gamma - (16\pi + 3\nu)\omega}{3\nu L} \quad (30)$$

Minimising respect to $\Phi$ we obtain:

$$\Phi_0 = 0 \quad \Phi_0 = \sqrt{\frac{-U_2}{2U_4}} \tag{31}$$

Equation 31 is used in Fig. 1E and Fig. 3 (dashed lines).

#### Estimation of the parameters in the theory

The mean spindle length and spindle volume are $2L = 57.1 \pm 0.7$ $\mu$m, $V_0 = (3.0 \pm 0.2) \times 10^4$ $\mu$m$^3$ ($n = 364$, mean $\pm$ SEM), where the volume was calculated by considering a tactoidal shape with the measured shape parameters (see Materials and Methods). Using standard error propagation we obtain the dimensionless volume $\nu \equiv V_0/L^3 = 1.27 \pm 0.08$. We consider a sufficiently small surface tension at the spindle poles such that in the absence of active dipolar stresses the spindle is found in an unfocused configuration $\omega/\gamma < \omega^\star/\gamma \simeq 0.5$ (see Fig. 1D). In order to deform the structure, contractility needs to be comparable to the surface tension at the walls and thus we expect $\sigma_c \sim \gamma/L$ as shown in Fig. 1D. Using the values $\sigma \simeq 70$ Pa and $\gamma \simeq 120$ pN/$\mu$m from Ref. (*4*), we obtain $\sigma L/\gamma \simeq 20$. For the poles to be almost completely focused with the previous contractility value, nematic elasticity needs to be $K/\gamma L \simeq 0.1$ in our model, assuming the one-constant approximation $K \equiv K_1 = K_3$. Deviations from this approximation do not change the results significantly (data not shown). Using the value for the surface tension $\gamma$, we obtain $K \simeq 340$ pN. Microrheology studies on spindles suggest that the Young's elastic modulus is found on the order of kilopascals (*5*). An estimation for the compressibility modulus of $A \sim 1$ kPa leads to a value $AL/\gamma \simeq 200$, consistent with the fact that the constant volume assumption is a very good approximation (see Fig. S2). Table 1 summarises all the main parameters used.

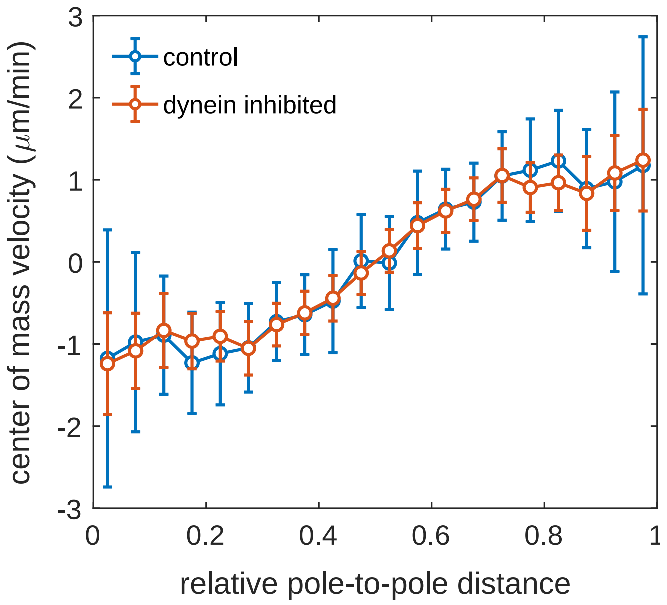


Figure S1: Center of mass microtubule velocity profile in control ($n = 13$) and dynein-inhibited spindles ($n = 10$) using fluorescent speckle microscopy data. Mean $\pm$ SD.

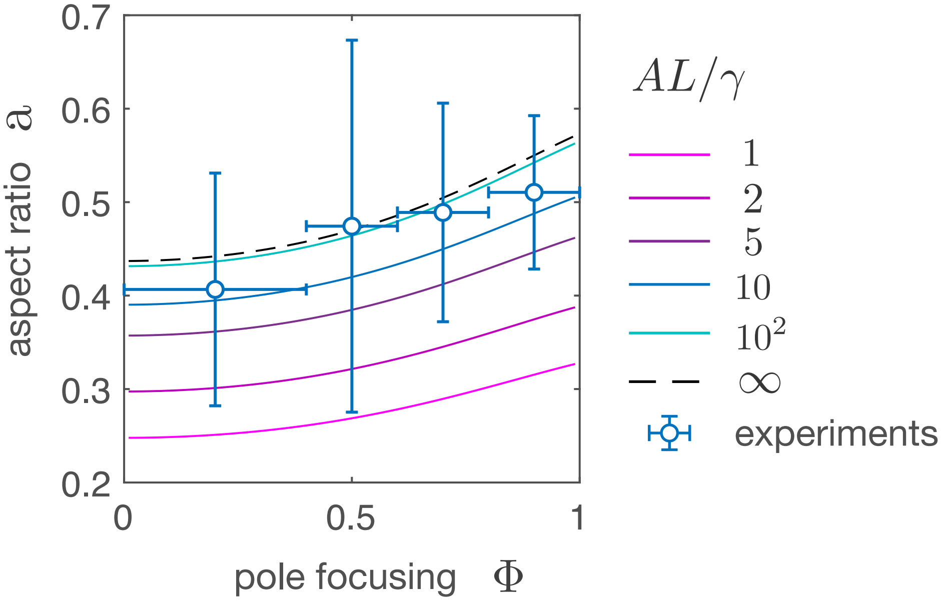


Figure S2: Curves in shape space for different values of the compression modulus $A$. The case $AL/\gamma \to \infty$ corresponds to the limit of constant volume. The experimental data is the same as in the inset of Fig. 3 in the Main Text. $K_1/K_3 = 1$, $K_1/\gamma L = 0.1$, $\omega/\gamma = 0.3$, $\nu = 1.2$.

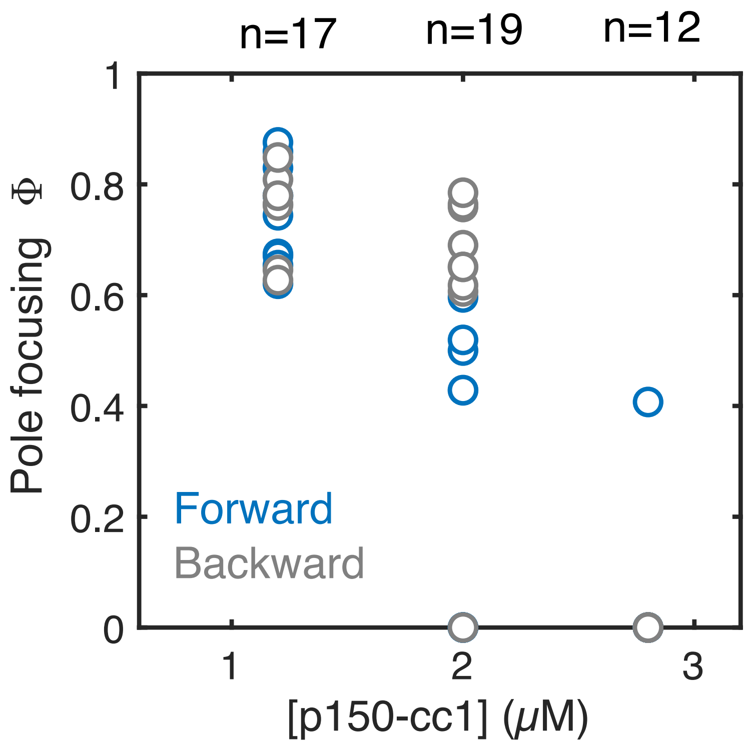


Figure S3: Example of a forward and backward titration during the same extract day showing the reversibility of the pole focusing process. Multiple circles from forward and backward titrations overlap at pole focusing $\Phi = 0$.

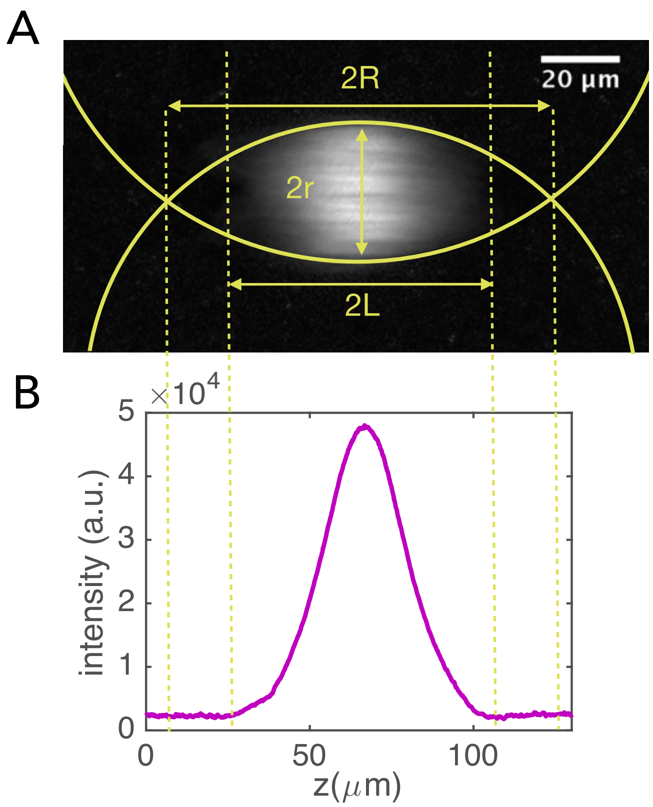


Figure S4: Shape characterization. A) Typical LC-Polscope retardance of a spindle with partially focused poles. The three diferent parameters $(R, r, L)$ were measured as indicated. B) Summed retardance intensity along the spindle short-axis. The threshold retardance value to define $2L$ is the background level.

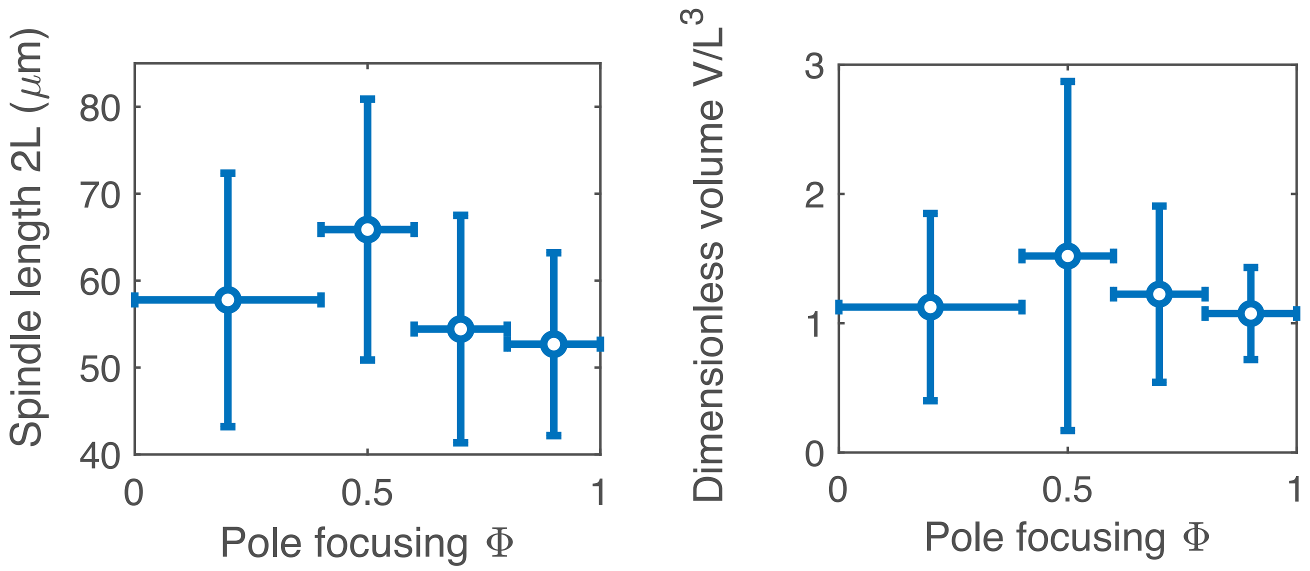


Figure S5: Spindle length $2L$ (left) and dimensionless spindle volume $V/L^3$ (right) as a function of the pole focusing parameter (mean $\pm$ SD, $n = 364$ spindles).

Table S1: Summary of the different parameters in the theory

| Parameter | Symbol | Value | Comment |
|---|---|---|---|
| Spindle half-length | $L$ | $28 \pm 7$ $\mu$m (mean $\pm$ SD) | exp. measure |
| Spindle half-width | $r$ | $13 \pm 5$ $\mu$m (mean $\pm$ SD) | exp. measure |
| Spindle volume | $V_0$ | $(30 \pm 2) \times 10^3$ $\mu m^3$ (mean $\pm$ SEM) | exp. measure |
| Dimensionless volume | $\nu$ | $1.27 \pm 0.08$ (mean $\pm$ SEM) | exp. measure |
| Dipolar contractile stresses | $\sigma$ | $\simeq 70$ Pa | Ref. (*4*) |
| Surface tension spindle wall | $\gamma$ | $\simeq 120$ pN/$\mu$m | Ref. (*4*) |
| Surface tension spindle poles | $\omega$ | $\lesssim 60$ pN/$\mu$m | estimation |
| Nematic elastic constant | $K$ | $\simeq 340$ pN | estimation |
| Compressibility modulus | $A$ | $\simeq 1$ kPa | estimation and Ref. (*5*) |
